# Regional associations of sleep architecture and Alzheimer’s disease pathology

**DOI:** 10.1101/2024.10.10.617528

**Authors:** Antonia Buchal, David Elmenhorst, Elena Doering, Verena Dzialas, Kathrin Giehl, Gérard N Bischof, Thilo van Eimeren, Alexander Drzezga, Merle C. Hoenig, the T-POT study group

**Affiliations:** University of Cologne, Faculty of Medicine and University Hospital Cologne, Department of Nuclear Medicine, Cologne, Germany; Research Centre Juelich, Institute for Neuroscience and Medicine II, Molecular Organization of the Brain, Juelich, Germany; German Centre for Neurodegenerative Diseases, Bonn/Cologne, Germany; University of Cologne, Faculty of Mathematics and Natural Sciences, Cologne, Germany; University of Cologne, Faculty of Medicine and University Hospital Cologne, Department of Neurology, Cologne, Germany

**Author notes:** **Corresponding Author:** Merle C. Hoenig, PhD, Institute for Neuroscience and Medicine II, Molecular Organization of the Brain, Research Centre Juelich; Wilhelm-Johnen-Straße 1, 52428 Jülich, Germany. Neuroimaging data used in preparation of this article were obtained from the T-POT study. As such, the investigators within the T-POT study group contributed to the design and implementation of T-POT, but did not participate in the analysis or writing of this report. A complete listing of T-POT investigators can be found in the acknowledgements.

## Abstract

**Objective:** Recent evidence suggests that disturbances of sleep architecture are linked to Alzheimer’s disease (AD) pathology. Here, we assessed the association between sleep architecture and regional amyloid and tau pathology employing a portable sleep-monitoring device in addition to PET imaging.

**Methods:** 18 cognitively normal adults (CN; M(Age) = 64.06 (8.63), Sex (M/F) =6/12) and 18 patients with MCI/early AD (M(Age) = 67.33 (8.25), Sex (M/F) =9/9) were included from the “Tau Propagation Over Time” (T-POT) study. All subjects underwent amyloid ([11C]-PiB) and tau ([18F]-AV1451) PET imaging. PET images were normalized to MNI-space and intensity standardized to the whole cerebellum ([11C]-PiB) or the inferior cerebellum ([18F]-AV1451). Sleep monitoring was performed at home using the portable “Dreem” EEG-headband (Beacon Biosignal), which is a reliable and comfortable wireless alternative to the gold-standard polysomnography (PSG). Sleep recordings were performed within six months of the PET acquisitions. At least, one sufficient night had to be acquired, which was used to assess the sleep macrostructure for each individual. Total duration of sleep phases per minutes (i.e. REM, N1, N2, N3) and total sleep time were extracted. In a first step, a linear mixed model (LMM) was used to compare the groups in terms of duration of the different sleep stages across the nightly recordings. Given the results of this comparison, mean N1 and N3 duration were subsequently correlated with regional amyloid and tau pathology SUVRs of 34 cortical regions using Spearman’s correlation. The reported results are based on one-tailed tests. All analyses were corrected for age.

**Results:** Patients with MCI/AD showed reduced N3 duration (*p* = .007) compared to the CN group. A trend was observed indicating that patients with MCI/AD exhibited longer N1 durations (*p* = .094); however, this difference did not reach statistical significance. Shorter N3 duration was associated with higher regional amyloid load in the paracentral lobe and the posterior cingulate gyrus, whereas longer N1 duration was linked to higher amyloid pathology in several regions, including the medial temporal lobe, cingulate cortex and the occipital lobe. Moreover, associations were observed between longer N1 duration and greater tau burden in regions comprising the temporal lobe, cingulate cortex, and medial-frontal areas of the brain.

**Conclusion:** Differences in sleep architecture between healthy controls and MCI/AD may arise from regionally-specific accumulation patterns of AD pathologies. Although it remains unknown whether disruptions in sleep architecture are a cause or a consequence, a complex relationship between AD-aggregation pathology in specific brain regions and the different phases of sleep appears to emerge.

## Introduction

The dementia prevention commission of Livingston et al. (2024) has recently reported that approximately 50 percent of worldwide dementia cases could be prevented by eliminating 14 risk factors. Interestingly, sleep is currently not listed among these modifiable risk factors, but a growing body of evidence suggests a significant association between sleep disturbances and risk of Alzheimer’s disease (see Rev. Mander, 2020). In this regard, shorter sleep duration and sleep disturbances, such as sleep apnea, have been associated with greater dementia risk (Sabia et al., 2021). In AD, it was reported that patients often suffer from sleep disturbances, such as a more fragmented sleep (Lim et al., 2013), less deep sleep (Sprecher et al., 2016), and less time spent in rapid-eye-movement (REM) sleep phases in comparison to healthy older individuals (Falgàs et al., 2023). Importantly, current findings suggest that disruptions in the sleep-wake cycle contribute to the build-up of the characteristic neuropathological hallmarks of AD (Roh et al., 2012), namely amyloid and tau pathology. Especially changes in slow-wave sleep (SWS) appear relevant in this regard (Mander et al., 2015; Winer et al., 2019), which may be due to the affectation of the so-called glymphatic system (see Rev. Nedergaard & Goldman, 2020). This system has recently been described to play a crucial role in clearing noxious proteins from the brain, in particular during deep sleep, through an exchange of the cerebrospinal with the interstitial fluid (Hauglund et al., 2020). Disruption of this system presumably results in reduced clearance and subsequent aggregation of neurotoxic proteins, such as amyloid β (Aβ) and tau tangles (Harrison et al., 2020; Hauglund et al., 2020). Notably, accumulation of these proteinopathies have recently been hypothesized to be linked to distinct changes in regional sleep activity patterns in the brain (see Rev. Mander, 2020). First studies reported on a significant association between amyloid pathology and modified SWS in the medial prefrontal cortex (Mander et al., 2015; Winer et al., 2020). In terms of tau pathology, it has been shown that lower SWS is associated with increasing tau pathology in orbitofrontal, entorhinal, parahippocampal, lingual and inferior parietal regions (Lucey et al., 2019; Winer et al., 2019). These associations may overall be of bi-directional nature. Thus, impairments in sleep may lead to increased pathology, whereby increased pathology may, in turn, lead to more sleep disturbances.

Notably, the current investigations on in vivo pathology and the macrostructure of sleep (i.e. different sleep phases) in AD have been conducted by using structured questionnaires on sleep behaviour or the gold standard of sleep assessment, namely polysomnography (PSG) (Carnicelli et al., 2019; Hita-Yañez et al., 2013). PSG comprises electroencephalography (EEG), electromyography (EMG), electrooculography (EOG) and measurement of respiratory activity (Arnal et al., 2020a) and is thus limited by its expensive, complicated and time-consuming nature. Given the recent introduction of portable EEG-sleep-monitoring headbands, sleep can nowadays easily be assessed at the homes of participants with low logistic effort and cost. Such measurements at the homes of the participants are more comfortable and thus facilitate sleep quality in a real-world environment (Kwon et al., 2021, Winer et al., 2019). Additionally, these portable devices may help avoid first night effects, which reflect differences from the initial night of monitoring due to the unfamiliar laboratory setting (Kwon et al., 2021; Wilson et al., 2022).

Leveraging on the advantages of the new portable devices, in this study we used a portable EEG-sleep-monitoring headband in a cohort of patients with mild cognitive impairment (MCI) or early AD and healthy controls to study differences in sleep macrostructure and their association with neuropathology using PET imaging data. We hypothesized that 1) differences in sleep macrostructure would be observable between healthy controls and MCI/AD patients, in particular in phases of SWS (e.g. N3 phase); 2) regions known for the regulation of sleep macrostructure (e.g. posterior cingulate cortex for regulation of SWS) would be related to regional AD pathology. Specifically, we assumed that increased pathology in these regions would be associated with a decrease in deep sleep phase duration (N3) and an increase in the duration of N1, the lightest phase of sleep.

## Materials and Methods

### Participants

In the current study, 36 participants (M(Age) = 65.69 (8.49), Sex (M/F) =15/21) from the ongoing “Tau Propagation Over Time” (T-POT) study were included. The T-POT study is funded by the German Research Foundation and aims to characterize the progression of AD and functional impairments at different stages of the disease. All participants were recruited from the University Hospital of Cologne. Main inclusion criteria for the T-POT study are: 1) no radiotherapy in the last 10 years, 2) absence of psychiatric diseases or claustrophobia, 3) age between 50-80 years of age, 4) no MRI contraindication. The entire T-POT study cohort is divided into four different groups according to established recommendation criteria by the NIA-AA (McKhann et al., 2011). Cognitively normal (CN) adults, who did not suffer from measurable objective or subjective cognitive decline; Patients with subjective cognitive decline (SCD), who reported self-perceived but not clinically measurable cognitive impairments; Patients with mild cognitive impairment (MCI) and patients with AD dementia. As part of the T-POT study, all participants undergo amyloid and tau PET imaging at baseline and 18-months follow-up. A sub-cohort of the T-POT participants were offered to participate in a study on sleep depending on the date of their PET scan acquisition and their willingness to participate in the sub-study.

For the current analysis of this study, individuals from this sub-cohort were selected based on the following criteria: 1) At least one night with a sleep recording quality greater than 80% (Arnal et al., 2020); 2) total sleep time greater than 240 minutes as expressed by an absolute z-score greater than or equal to three; 3) PET scan acquisition within 6 months of sleep recording. Due to the small portion of SCD patients with available sleep data, this group has been excluded from the current analysis. These criteria resulted in 18 CN, 15 MCI and 3 AD patients. The MCI and AD group was clustered into one group given the relatively small sample size (see Table 1 for an overview of participant characteristics). The study was approved by the Ethics Committee of the University of Cologne and all procedures were performed according to the Declaration of Helsinki. All participants provided written informed consent.

**Table 1.** Group characteristics. Means and standard deviations for the group demographics are presented. Chi-square test was used to test for differences in sex. Independent two-tailed t-tests were conducted to determine differences in continuous variables like age, education, amyloid burden and tau load; *p < .05, **p < .001. MMSE = mini mental state examination; SUVR = standard uptake value ratio; ROI = region-of -interest.

|  | **CN**<br>(n = 18) | **MCI/AD**<br>(n = 18) | **CN vs. MCI/AD** |
| --- | --- | --- | --- |
| Age (years) | 64.06 (8.63) | 67.33 (8.25) | $t(34) = -1.17, p = .252$ |
| Sex (M/F) | 6/12 | 9/9 | $X^2(1) = 1.00, p = .317$ |
| MMSE (points) | 29.44 (0.51) | 27.33 (2.45) | $t(34) = 3.58, p = .001^*$ |
| Education (years) | 17.25 (1.98) | 15.38 (3.75) | $t(34) = 1.06, p = .296$ |
| Global amyloid (SUVR) | 1.07 (0.21) | 1.55 (0.48) | $t(34) = -3.95, p < .001^{**}$ |
| Tau in meta-ROI (SUVR) | 1.18 (0.10) | 1.52 (0.47) | $t(34) = -2.95, p = .006^*$ |

### PET Imaging

The Siemens Hybrid-Scanner 3T MR-PET with a high-resolution BrainPET and a 3T MRT MAGNETOM Trio was used to acquire all imaging data. All participants underwent dynamic scanning for at least 70 and 100 minutes to obtain ^11^C-PIB and ^18^F-AV1451 scans. All scans were averaged across specific timeframes and co-registered to the corresponding MRI image. Spatial normalization was performed on all scans using statistical parametric mapping (SPM12; https://fil.ion.ucl.ac.uk/spm). Standardized uptake value ratios (SUVR) were calculated for every scan with reference to each individual’s cerebellum. Next, mean SUVRs were extracted using a global cortical mask for the amyloid PET scans and an established meta region-of-interest (ROI) (Jack et al., 2017) including the medial temporal lobe for the tau PET scans. Moreover, mean SUVRs for each participant were extracted based on the pre-processed amyloid and tau PET scans for 34 cortical regions according to the Desikan parcellation scheme (Desikan et al., 2006) and averaged across both hemispheres. The averaging was performed to account for the anticipated symmetry of pathology across brain regions and to reduce the number of comparisons between sleep phases and regional pathology load.

### Measurement of sleep macrostructure

Sleep architecture was assessed using the sleep-monitoring headband (SH) by Beacon Biosignals (https://beacon.bio/dreem-headband/), which is a wireless device that records and stores physiological sleep data throughout a full night of sleep. All recordings were performed at home which can reflect real-life sleep. Two sensors types embedded in the SH can measure various sleep-related physiological signals: five dry EEG electrodes measure cortical brain activity, achieving seven derivations (FpZ-O1, FpZ-O2, FpZ-F7, F8-F7, F7-O1, F8-O2, FpZ-F8; sampled with 250 Hz with a 0.4–35 Hz bandpass filter). The 3D accelerometer positioned over the head captures movements, positions and the breathing frequency. Sleep physiology can then be extracted using the automatically applied state-of-the-art machine learning algorithm provided by Beacon Biosignal.

Participants were asked to wear the SH for a minimum of three consecutive nights. To investigate sleep macrostructure, total sleep time (TST) was obtained for each subject for the respective nights. Beyond that, the duration (in minutes) of N1, N2, N3 and REM sleep were extracted for each night and each participant and submitted to the analyses.

### Statistical analysis

#### Linear mixed model analysis for group comparisons of sleep phases

Using a linear mixed effects model (LMM) without random effects and a scaled-identity matrix structure, the two groups were compared in terms total sleep time and the actual duration of time spent in N1, N2, N3 and REM sleep. For every patient, four full night recordings were considered as repeated measures. Recordings for night five and six were excluded due to insufficient variance caused by a decreasing number of measures across individuals. The model was set up considering age as a covariate in addition to group as fixed effect. Adjustment for multiple comparisons was performed using the Bonferroni correction (q<0.01).

#### Correlation analyses between AD pathology and sleep disturbances

Using SPSS, non-parametric one-tailed Spearman correlations were performed to evaluate the association between the duration of time spent in different sleep phases and regional AD pathology in the 34 cortical regions. To do so, the averaged regional mean SUVR of amyloid and tau as well as averaged N1 and N3 duration across the 4 nights were used, respectively. Given that N1 and N3 are assumed to be modified in AD (Targa et al., 2021), we specifically focused on the expression of these sleep phases. This was additionally done to reduce the number of regional correlations.

## Results

### Linear Mixed Model Analysis for group comparisons of sleep phases

The LMM did not show significant differences in terms of total sleep time (ßTST = 10.42, CI = -19.74:40.58, *p* = .495). However, regarding sleep macrostructure, a in significant differences in N3 duration was observed (ßN3 = 16.95, CI = 4.82:29.07, *p* = .007). Additionally, a trend was found in N1 duration (ßN1 = -3.73, CI = -8.11:0.64, *p* = .094), although this result did not reach statistical significance. No significant effect of age was indicated. The MCI group demonstrated lower N3 (Mean = 65.50 min) and higher N1 (Mean = 34.63 min) duration in comparison to the CN group (N3: Mean = 82.30 min, N1: Mean = 31.81 min) across the four nightly measurements, respectively. No significant differences were found in N2 (ßN2 = 9.01, CI = - 13.31:31.16, *p* = .422) and REM (ßREM = -11.80, CI = -33.82:10.22, *p* = .291) duration. After multiple comparison correction, N3 duration remained significantly different between groups. Figure 1 illustrates the differences in sleep phases between groups, represented by the average duration of distinct sleep phases for CN and MCI/AD.

**Figure 1.**
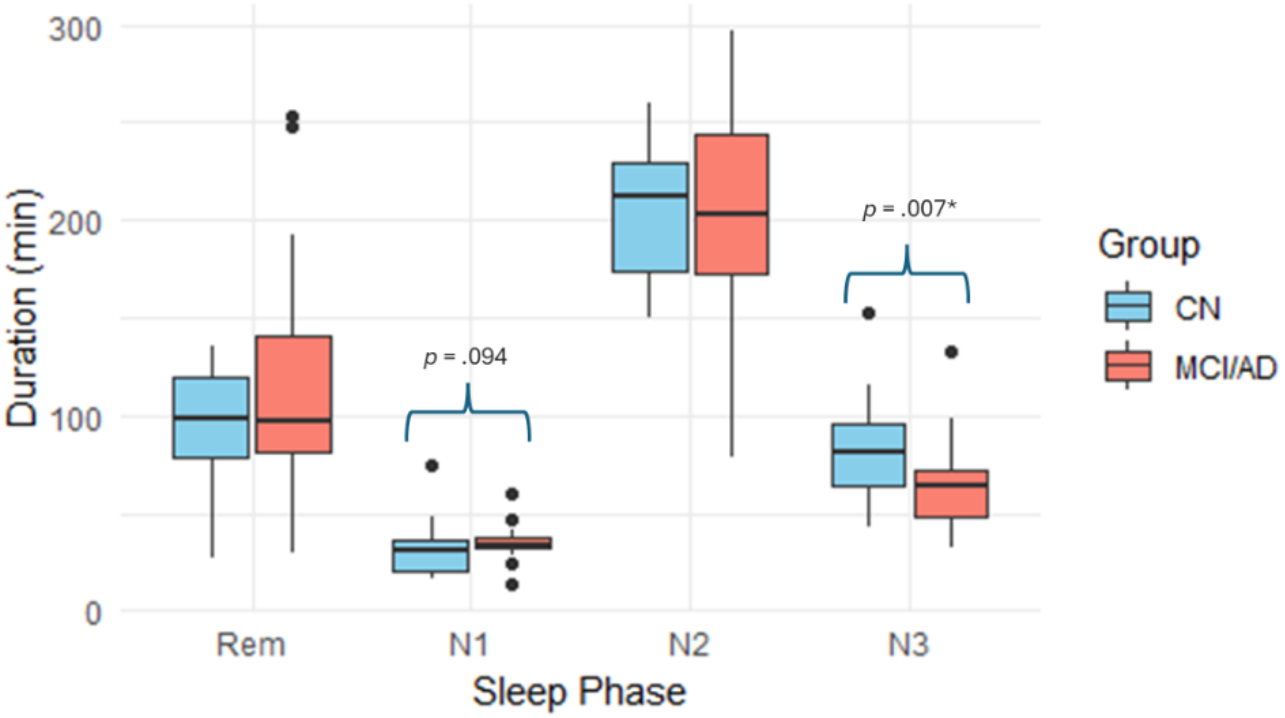
Group difference in the macrostructure of sleep. Average duration in minutes of each sleep phase across the four nights for each group is depicted. Groups are color-coded and (trend) significant differences are highlighted.

### Correlation analysis between Alzheimer’s disease pathology and sleep macrostructure

#### Amyloid pathology

Significant associations were observed for regional amyloid and tau pathology with N3 and N1 duration. In particular, there was a significant negative correlation between N3 duration and amyloid pathology within the paracentral lobe (*r* = -.29, *p* = .05) and trend significant association with the posterior cingulate cortex (*r* = -.27, *p* = .06).

Positive associations were found between N1 and amyloid burden in the following regions: caudal anterior cingulate cortex (*r* = .29, *p* = .05), isthmus cingulate cortex (*r* = .30, *p* = .04), lingual gyrus (*r* = .41, *p* = .01), pericalcarine cortex (*r* = .30, *p* = .04) and in the parahippocampal gyrus (*r* = .37, *p* = .01).

#### Tau pathology

No significant correlations were found between N3 and regional tau load. For tau and N1 duration, the following regions demonstrated a significant positive effect: bank of superior temporal sulcus (*r* = .30, *p* = .04), entorhinal cortex (*r* = .32, *p* = .03), fusiform gyrus (*r* = .32, *p* = .03), inferior temporal gyrus (*r* = .30, *p* = .04), isthmus cingulate (*r* = .32, *p* = .03), lingual gyrus (*r* = .30, *p* = .04), parahippocampal gyrus (*r* = .37, *p* = .01), pars opercularis (*r* = .28, *p* = .05), posterior cingulate cortex (*r* = .28, *p* = .05) and precentral gyrus (*r* = .30, *p* = .04).

All p-values reported reflect one-tailed tests. In Figure 2, the respective regional associations with N1/N3 duration for amyloid and tau pathology are depicted.

**Figure 2.**
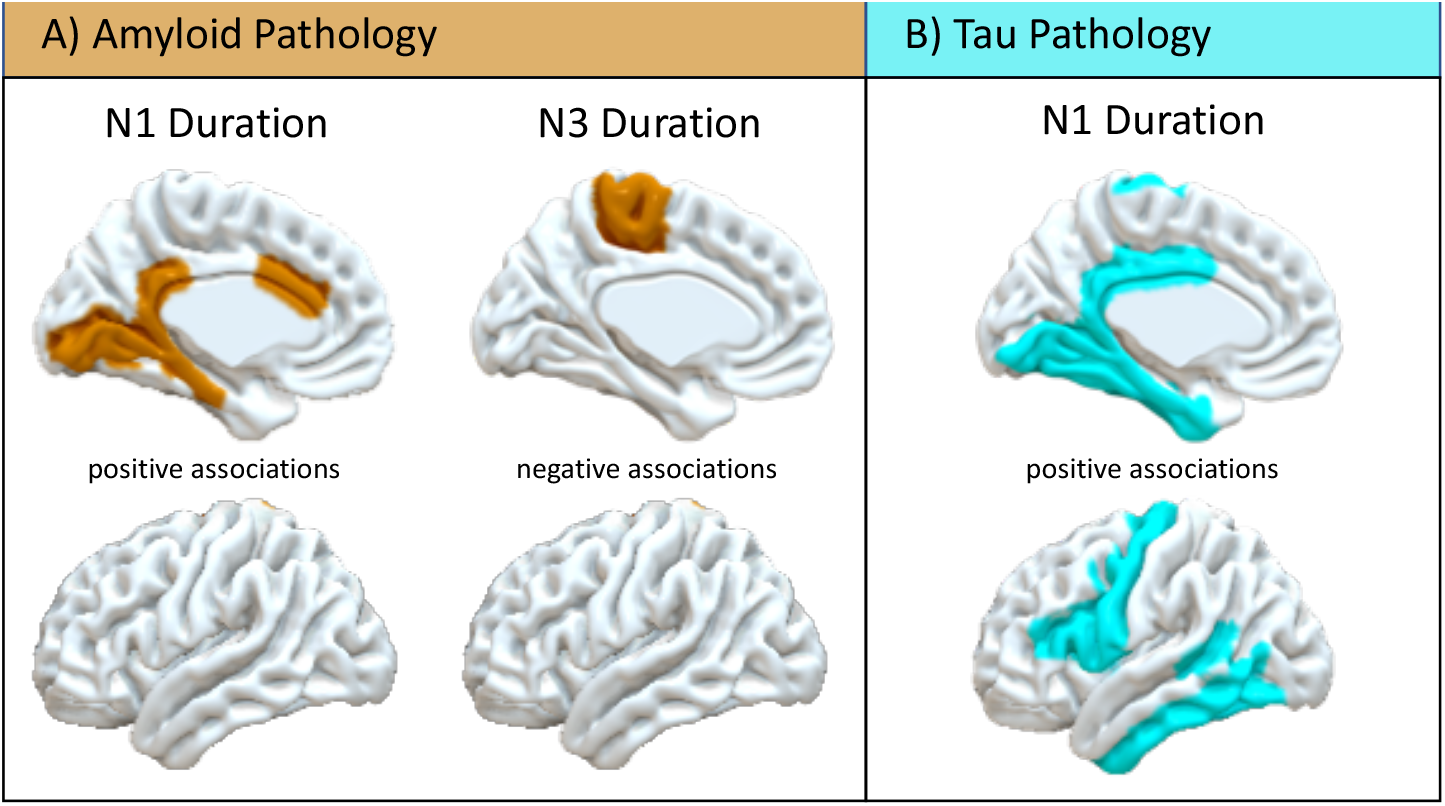
Atlas regions that showed associations of amyloid and tau pathology with N1 and N3 duration. A) Positive associations between local amyloid pathology burden and N1 and negative associations with N3 duration are depicted. B) Positive associations of local tau pathology and N1 duration are presented. No findings were found in terms of tau pathology and N3 duration.

## Discussion

In this study, we aimed to investigate changes in sleep macrostructure linked to regional AD pathology as assessed by PET imaging. Using a portable EEG-sleep-monitoring device in MCI/AD patients, we were able to show reductions in N3 duration and a concomitant increase in N1 duration in the MCI/AD group vs. healthy controls. Shorter N3 duration was associated with increased amyloid pathology in the paracentral lobe and posterior cingulate cortex, while a longer N1 duration correlated with regional amyloid pathology in the caudal anterior cingulate cortex and temporal regions. In terms of tau pathology, significant positive associations of N1 were observed with tau pathology in hippocampal, temporal, posterior cingulate, and precentral regions. The results indicate that local changes in sleep architecture may potentially arise from regionally-specific accumulation patterns of AD pathologies.

Notably, the observation of N1 and N3 duration being different between MCI/AD and cognitively normal controls is in line with previous finding using traditional PSG (Ju et al., 2017; Targa et al., 2021). Indeed, current assumption is that increased N1 compensates for decreasing N3 and lower slow wave activity during deep sleep (Ju et al., 2017). Importantly, several studies have demonstrated that changes in sleep macrostructure are linked to cognitive performance, such as memory consolidation (Backhaus et al., 2007; Miyata et al., 2013; Zhu et al., 2012). The observed characteristics of sleep macrostructure in the MCI/AD group may thus partly have contributed to lower cognitive performance level in this group. Presumably, these associations between sleep and cognitive performance may be mediated or driven by the underlying neuropathological burden in terms of amyloid plaques and tau tangles.

To gain a more comprehensive understanding on the regional interplay between neuropathology and sleep architecture, we therefore associated regional amyloid and tau pathology with the respective sleep phase duration of light (N1) and deep sleep (N3). We thereby identified sleep-pathology patterns, comprising regions known for the regulation of sleep macrostructure such as for example the anterior and posterior cingulate. The anterior cingulate cortex is responsible for suppressing constant waking (Atlan et al., 2024) and functioning has been reported to be impaired in AD (Jones et al., 2005), potentially by greater amyloid burden as signified in our study. Disruption of the function of the anterior cingulate cortex may, in turn, explain more fragmented sleep and why AD patients remain in N1 and enter less often deeper sleep stages. Moreover, we observed a negative association between N3 duration and amyloid pathology in the posterior cingulate cortex. This brain region is involved in generating slow waves during deep sleep (Murphy et al., 2009) and amyloid accumulation in this region has been hypothesized to lead to local deficits in non-REM sleep (see Rev. Mander, 2020), as indicated by our current results. Regarding tau pathology, a positive association between N1 and tau accumulation within the medial temporal lobe (MTL) was observed, which also includes the entorhinal cortex and the parahippocampal gyrus. This observation is line with recent findings suggesting local tau pathology in the parahippocampal gyrus and other temporal and frontal regions being associated with reduced neural activity and sleep disturbances in patients with early AD (Wang & Peng, 2021). Interestingly, according to the authors, patients developed sleep disturbances after the onset of cognitive symptoms, indicating that the same pathological processes may be conductive for modification of sleep macrostructure and that sleep disturbances may be a consequence of AD. Notably, regions affected by both, amyloid and tau pathology, such as the isthmus cingulate, lingual gyrus and parahippocampal gyrus, are associated with a prolonged N1 duration in MCI/AD patients compared to healthy adults. This may also suggest that disrupted sleep patterns may be a consequence of the disease, particularly in later stages of the disease, when the formation of neurofibrillary tangles is already triggered by amyloid, as suggested by the amyloid cascade hypothesis by Hardy & Higgins (1992).

A few limitations need to be considered in the interpretation of the current results. First, the rather small sample size warrants replication in larger cohorts to increase effect size and power of the analyses. Moreover, correlations were tested in a one-tailed manner instead of the conventional two-tailed approach and were not corrected for multiple comparisons, which may have led to an increase in alpha-error. Moreover, given that we used data of MCI and early AD patients, AD pathology, in particular, amyloid pathology may already have distributed throughout the brain increasing regional dependence in terms of pathology load. Nevertheless, we were able to determine distinct regions of pathology load to be linked with the length of N1 and N3 sleep phases. As these regions have previously been reported to be involved in sleep production, the current results support the notion of the potential bi-directional relationship between sleep and AD pathology.

Overall, the current findings using a novel wearable technology in combination with molecular imaging emphasize the interplay of local pathology burden and modified sleep macrostructure in cognitively impaired individuals. Further longitudinal studies in at risk populations of developing AD are required to test whether the associations between regional pathology and sleep changes are a cause or a consequence of the disease. Better understanding of the mechanistic pathways on how sleep changes are related to the pathology build-up in AD will provide important insights that will be relevant for the development and timely implementation of therapeutical interventions by the modulation of sleep.

## Funding

MCH received funding from the Alzheimer Forschung Initiative e.V. for the conduct of the sleep study. The neuroimaging data was acquired as part of the fully-funded T-POT study by the German Research Foundation (DR 445/9-1).

## Acknowledgements

The T-POT study group consists of several researchers from different disciplines, including medicine, neuropsychology, radiochemistry and physics. Significant contributors to the study are: Elena Jäger^1^, Andreas Matusch^2^, Tina Kroll^2^, Andreas Bauer^2^, Sabine Klein^2^, Stephanie Krause^2^, Philipp Krapf^3^, Bernd Neumaier^3^, Christoph Lerche^4^, Lutz Tellmann^4^, Silke French^4^, Philipp Zeyn^5^, Frederik Sand^5^, Alfredo Ramirez^5^, Frank Jessen^5^, Nils Richter^6^, Hendrik Theis^6^, Özgür Onur^6. 1^ University of Cologne, Faculty of Medicine and University Hospital Cologne, Department of Nuclear Medicine, Cologne, Germany; ^2^ Research Centre Juelich, Institute for Neuroscience and Medicine 2, Molecular Organization of the Brain, Juelich, Germany; ^3^ Research Centre Juelich, Institute for Neuroscience and Medicine 5, Radiochemistry, Juelich, Germany; ^4^ Research Centre Juelich, Institute for Neuroscience and Medicine 4, Medical Imaging Physics, Juelich, Germany; ^5^ University of Cologne, Faculty of Medicine and University Hospital Cologne, Department of Psychiatry, Cologne, Germany; ^6^ University of Cologne, Faculty of Medicine and University Hospital Cologne, Department of Neurology, Cologne, Germany.

## Conflict of Interest

The first author and co-authors report no conflicts of interest related to this work. MCH reports the provision of 15 sleep-monitoring headbands for scientific purposes by the company Dreem (now Beacon Biosignal) as part of a research proposal prize.

## Notes

### Competing Interest Statement

The authors have declared no competing interest.

